# Sequence data from the internal and external transcribed spacers of nuclear ribosomal DNA of *Cyclamen purpurascens* allow geographic mapping

**DOI:** 10.1101/432203

**Authors:** Hannah Miriam Jaag

## Abstract

*Cyclamen purpurascens* (Alpine, European or purple cyclamen) is native to central Europe. Since decades it is discussed wether the occurrences of *C. purpurascens* north of the alps is native or if it was introduced. Here the nuclear ribosomal DNAs (rDNA) are sequenced in oder to obtain a phylogenetic geographic pattern. Phylogenetic analyses of ITS and NTS/ETS sequences distinguish three main clades coinciding with geographical distribution: Eastern alps (Austria), southern alps (Switzerland, Italy) and western Alps (France). The paper presents interspecific relationship of C. purpurascens based on geographic sequences of rDNA. The observed variations suggest that some plants were introduced via Benedictine gardens and the plants from Monastery gardens seem to origin from Lower Austria.

## Introduction

*Cyclamen purpurascens* (Alpine, European or purple cyclamen) is a species in the genus *Cyclamen* of the family Myrsinaceae, formerly Primulaceae, and is native to central Europe from eastern France across the Alps to Slovakia and south to Croatia. North of the alps there are some occurrences of *C. purpurascens* like in southwest Germany (Mühlheim, Kisslegg, Salem, Brigachtal) and Switzerland (Schaffhausen) [1]. Welten & Sutter recorded the Jura to Oensingen (Mümliswil), Lake Lucerne and the Upper Lake Zurich as the next environs [2]. Since decades it is discussed wether the occurrences of C. purpurascens north of the alps is native or if it was introduced.

In 2011 Slovak *et al*. published an interesting paper about sequencing noncoding DNA regions of *C. purpurascens* to distinguish geographic origins [3]. Unfortunately the polymorphisms in the sequenced regions were too low to generate an useful mapping. Nevertheless the idea of geographic mapping by sequencing of specific DNA regions seems to be challenging but also fascinating. Since chloroplast DNA sequences don’t give enough data for phylogenetic analysis of C. purpurascens we investigated different approaches.

In an analysis of phylogenetic interrelationships of the genus Cyclamen L., Anderberg *et al*. sequenced the internal transcribed spacers of nuclear ribosomal DNA [4]. Ribosomal DNA (rDNA) consists of a tandem repeat of an operon, composed of non-transcribed spacer (NTS), external transcribed spacer (ETS), 18S, internal transcribed spacer 1 (ITS1), 5.8S, internal transcribed spacer 2 (ITS2) and 26S tracts (Fig 1). In preliminary studies we could find some variations not just in ITS1 and ITS2, but also upstream of 18S in the region of NTS/ETS in C. purpurascens from different geographic regions.

**Fig 1.**
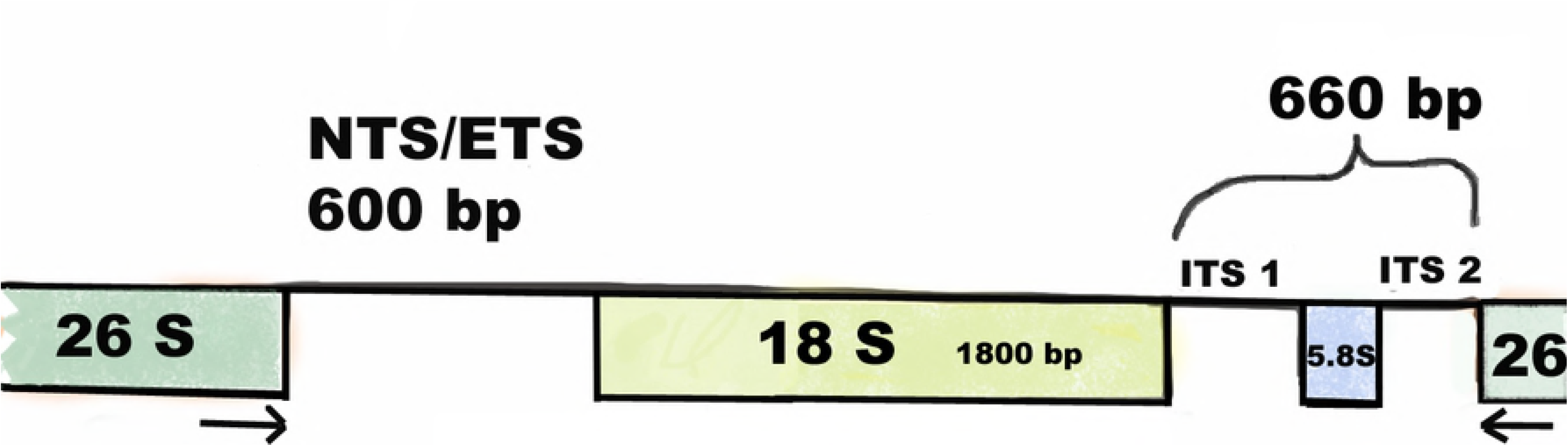
Schematic representation of ribosomal DNA (rDNA). *C. purpurascens rDNA of* NTS/ETS, 18S, ITS1, 5.8S and ITS2 is a part of a tandem repetitive cluster of 3155 basepairs. The rDNA contains NTS/ETS of ∼600 basepairs, 18S of ∼1800 basepairs, ITS1/5.8S/ITS2 of ∼660 basepairs and 26S.

The aim of this study was to investigate the sequence of the NTS/ETS, ITS1 and ITS2 to do a geographic mapping of *C. purpurascens*. For this we collected and sequenced 97 leaves of alpine cyclamen of different geographical origin.

## Materials and Methods

### Plant material

A total of 97 leaves of *C. purpurascens* from different geographic origins were collected and rDNA was sequenced (Fig 2; Table 1).

**Fig 2.**
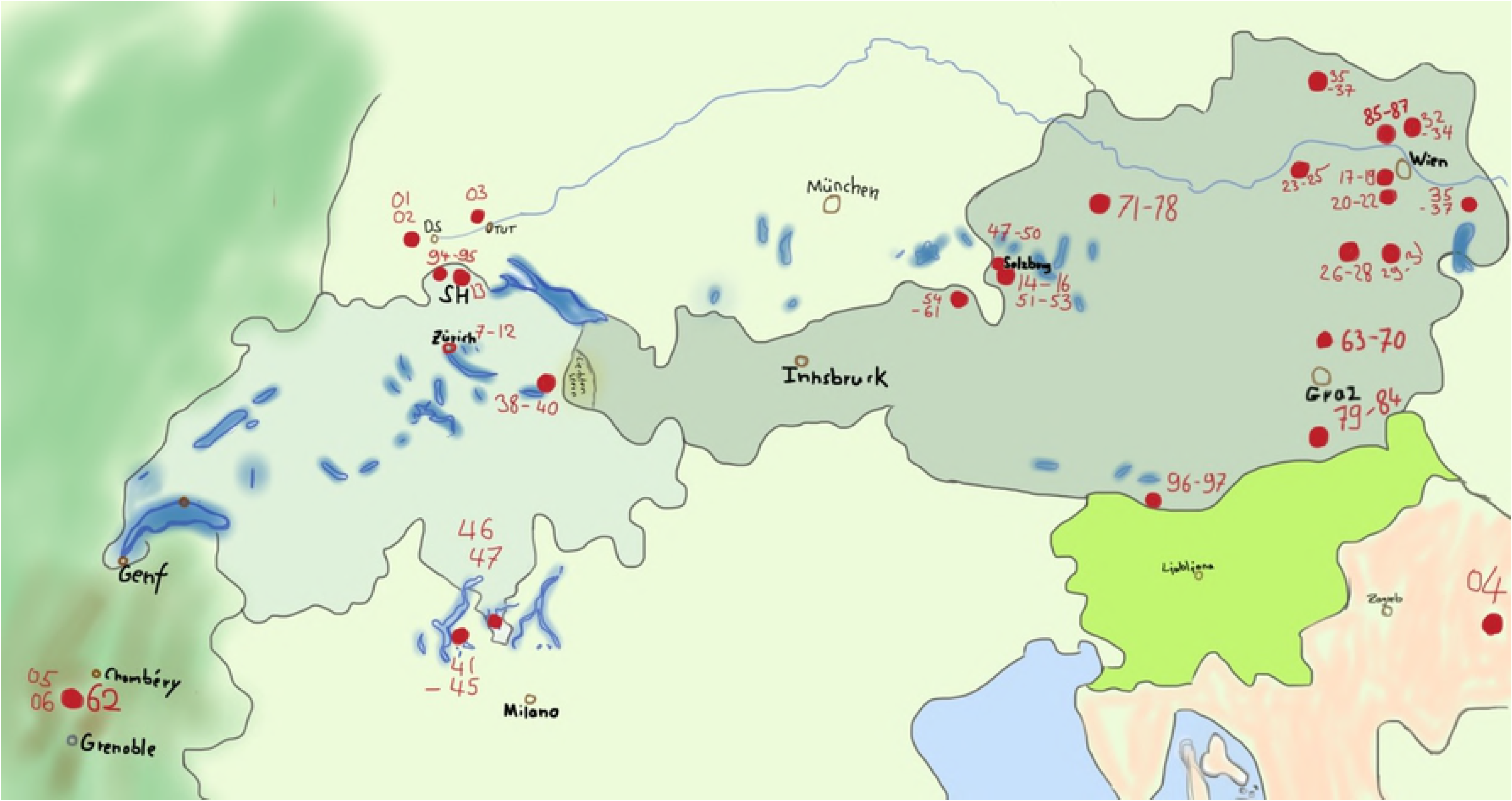
Map of central Europe. Analyzed samples an their voucher numbers are indicated in red.

**Table 1.**
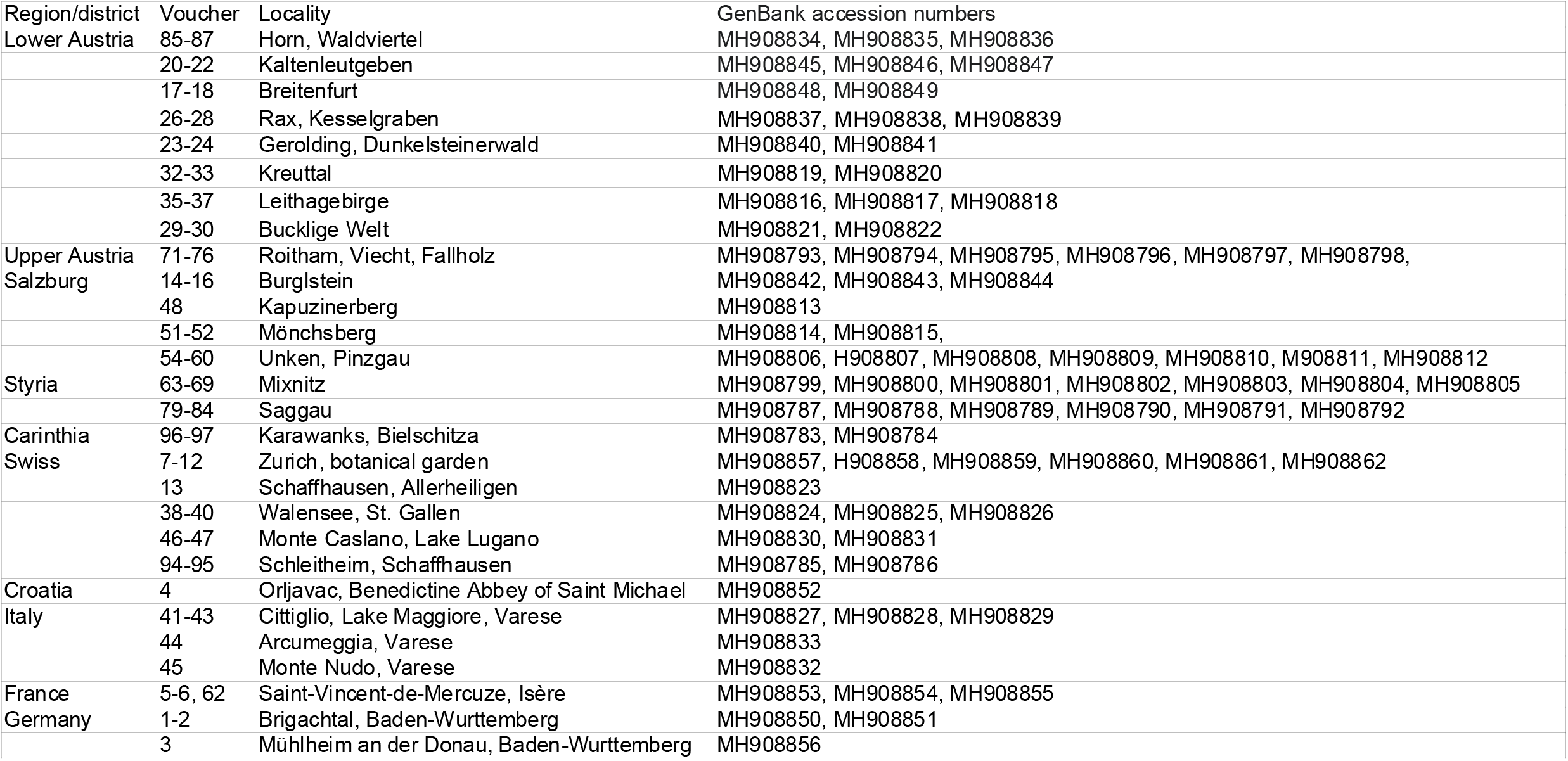
Plant materials. Plant materials used. Region, voucher information, localities and GenBank accession numbers are indicated.

### DNA region

DNA sequences from the ribosomal DNA were used for the present analysis. Ribosomal DNA (rDNA) consists of a tandem repeat of an operon, composed of non-transcribed spacer (NTS), external transcribed spacer (ETS), 18S, internal transcribed spacer 1 (ITS1), 5.8S, internal transcribed spacer 2 (ITS2) and 26S tracts. The spacers are regions within the nuclear ribosomal DNA gene that separates the 18S, 5.8 S and 26S genes. The ITS1 and ITS2 regions have been used in several phylogenetic reconstructions and proved to be variable to a suitable degree for investigations [4]. Here we also analyzed the NTS/ETS region (Fig 1).

### DNA extraction, PCR and sequencing

50 mg of leave material was grinded in 1 ml buffer HS (10 mM Tris-HCl, pH 7.6, 10 mM KCl, 10 mM MgCl2, 400 mM NaCl, 2 mM EDTA, 1% SDS, 0,1 mM DTT, 20 µg Proteinase K, 10 µg RNAse) and incubated for 3 hours at 50°C. Debris was collected by centrifugation and DNA was precipitated with isopropanol and washed with 70% EtOH. Genomic DNA was also purified with columns from Roti Prep Genomic DNA kit from Roth.

PCR was performed with Polymerases (One Taq or Phusion) from New England BioLabs and primers indicated in Table 2. Temperatures were calculated with Tm Calculator from New England BioLabs. NTS/ETS was amplified with primer HM95/HM96.18S was amplified with primer HM84/93, HM92/89 and HM94/93. ITS1/5.8S/ITS2 with primer HM102/HM82 or HM81/HM82. PCR products were gel purified with Monarch DNA Gel Extraction Kit from New England BioLabs and sequenced.

**Table 2.**
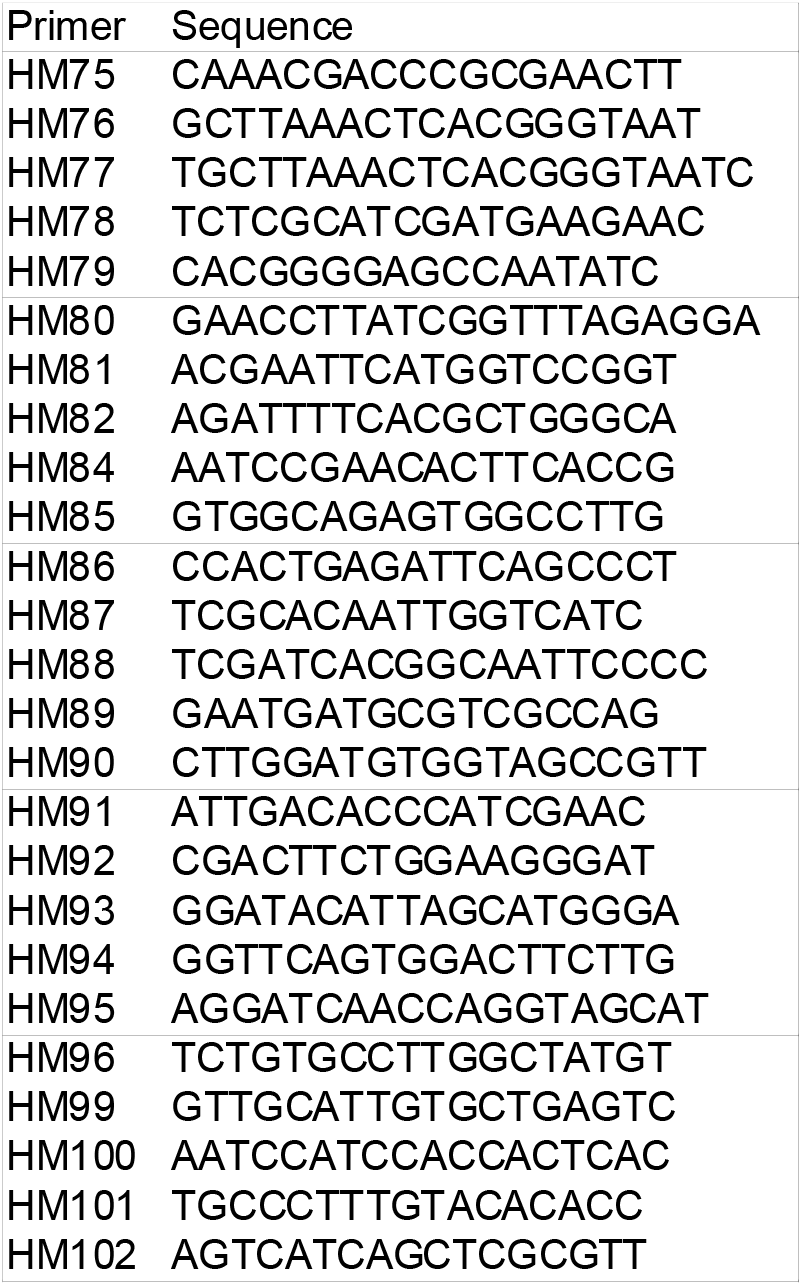
**Primers** Primers used for this study.

The PCR product of NTS/ETS was sequenced with primes HM96, HM100 or HM 95. The PCR product of 18 S was sequenced with primers HM 84, HM93, HM92, HM89, HM94. The PCR product of ITS1/5.8S/ITS2 was sequenced with primes HM78, HM82 or HM79. Sequencing was performed by GATC Biotech (now Eurofins Genomics) in Cologne. Sequences were analyzed by Multiple Sequence Alignment by CLUSTALW, Kyoto Bioinformatics Center and SeqDoC by the Bioinformatics Facility of the ARC Special Research Centre for Functional and Applied Genomics.

## Results

### Polygenetic analyses of *C. purpurascens* collected in different geographic regions

We sequenced 97 leaves of different geographical origin of C. purpurascens in the region of NTS/ETS and ITS1/5.8S/ITS2 (Table 1; Fig 1 and 2).

Multiple sequence alignment was done with the 3155 basepair rDNA region (Fig 1). For a better overview some similar sequences were omitted. Also five too divergent sequences were omitted. This were one sequence from Unken (Pinzgau), one from Salzburg, two from Styria and one from Arcumeggia (Italy). Plants from Switzerland (Walensee, St. Gallen and Monte Caslano, Lake Lugano, Schaffhausen) are related to the plants collected in Italy (region Lake Maggiore). Another cluster could be found with three plants from France (St. Vincent de Mercuse, Isère, North of Grenoble). In Austria plants get more diverse in the analyzed rDNA region, the farer we collect samples from Lower Austria (Fig 3). So plants from Styria (Mixnitz) and also from Pinzgau (Unken) are highly diverse. Surprisingly leaves collected in Lower and Upper Austria are very much conserved in the region of NTS/ETS/18S/ITS1/5.8S/ITS2. This conserved sequence could also be found in some plants from Salzburg, in a sample from Schaffhausen (Switzerland), Mühlheim an der Donau (Germany) and also from Orljavac (Croatia). The conserved sequence could not be found in the plants from Brigachtal (Germany) and from Schleitheim (Schaffhausen, Switzerland). The plants with the conserved sequence similar to the region of Vienna are all growing close to Benedictine gardens. So the leaf from Schaffhausen was collected in the garden of the former abby Allerheiligen, Mühlheim an der Donau is close to Beuron Archabbey and near Orljavac was a Benedictine abbey of Saint Michael.

**Fig 3.**
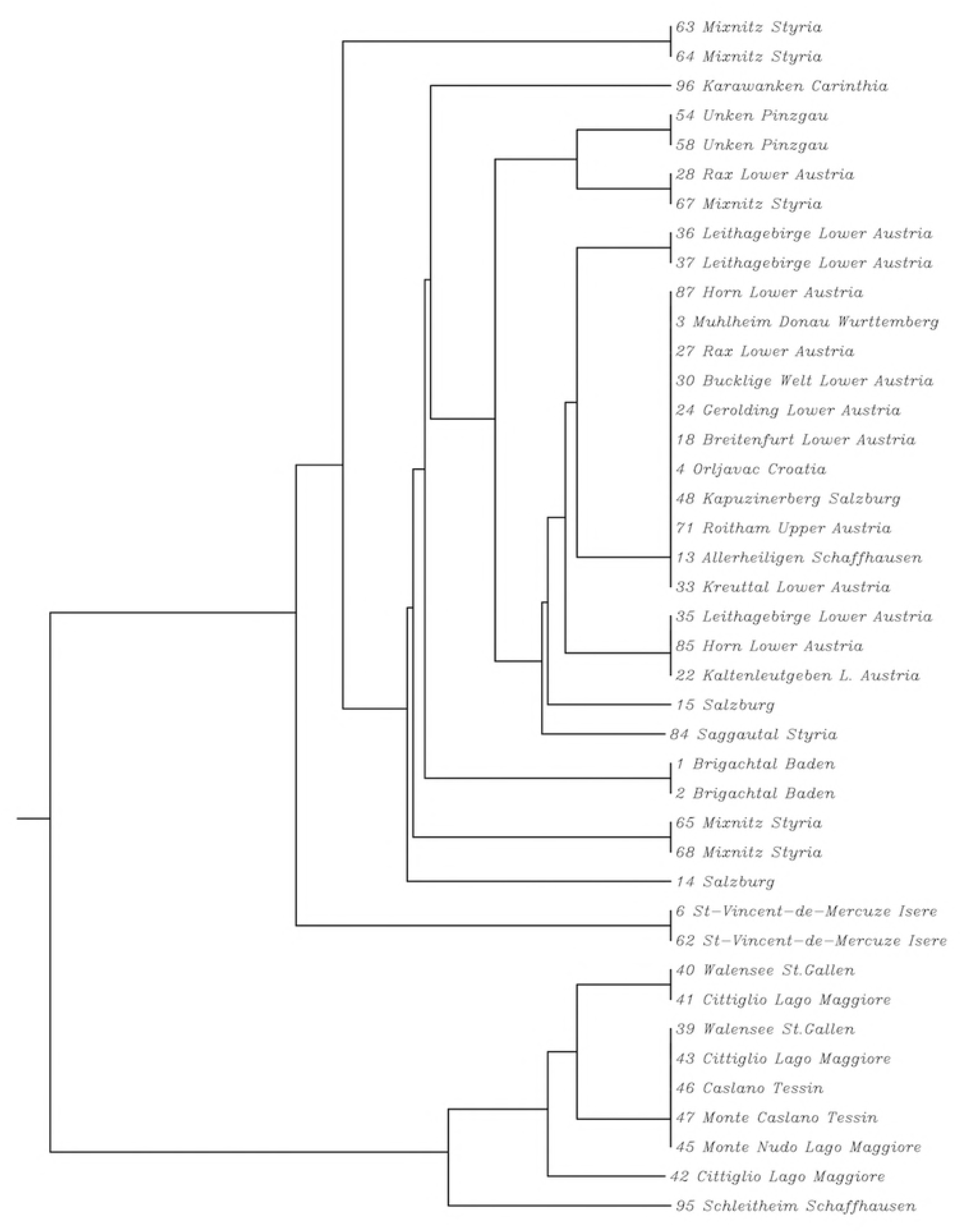
Polygenetic tree of *C. purpurascens* from different geographic origins. Plants from Switzerland (Walensee, Monte Caslano, Schleitheim) are related to the plants collected in Italy (Cittiglio, Caslano, Monte Nudo). Another cluster could be found with plants from France (St. Vincent de Mercuse). In Austria plants get more diverse in the analyzed rDNA region, the farer from Vienna. Plants from Styria (Mixnitz) and also from Pinzgau (Unken) are highly diverse. Leaves collected in Lower and Upper Austria are very much conserved in the region of ETS/18S/ITS1/5.8S/ITS2. This conserved sequence could also be found in some plants from Salzburg, in samples from Schaffhausen, Mühlheim and also from Orljavac. It could not be found in the plants from Brigachtal and Schleitheim.

### Sequence comparison of geographic regions

Sequence alignment comparison of 50 bases in the ETS of 14 plants from different origins show the difference of plants from Switzerland, Italy, France and Austria. So all analyzed plants from Austria and France have an additional G (TGGCCC) which is missing in plants from Italy and Switzerland (TGCCC). Nine bases downstream of this signal plants from France are having a unique signal of AAAA instead AAGA like the analyzed plants form Austria, Switzerland and Italy (Fig 4). Similar signals could be found in the ITS1 and ITS2 of the rDNA (data not shown).

**Fig 4.**
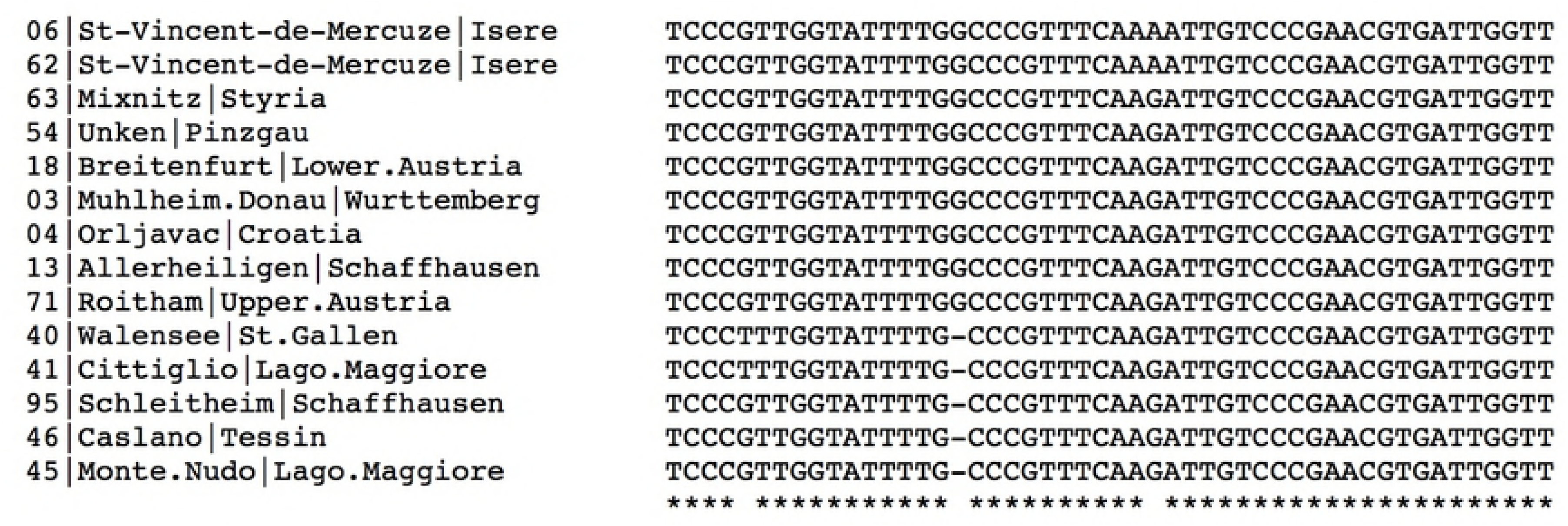
Sequence alignment of nt 100–150 in the NTS/ETS of selected plants. Plants from Italy and Switzerland show the signal TTGCC and plants from Austria and France TTGGCC. The plants from France can be distinguished 9 bases downstream by AAAA instead of AAGA in Austria, Italy and Switzerland.

### Sequence alignment of ETS from two plants of different origin

Downstream of this signals we could also find some single nucleotide differences in the region of ETS. Using two chromatograms, we aligned images of the two chromatograms. Here we compare nucleotide 100–500 of ETS of a leaf from Lower Austria to the corresponding sequence of a leaf from Cittiglio (Fig 5). The difference profile in the middle allows identification of base substitutions, insertions and deletions. As shown in Fig 5., at the 5’ are two G in the plant from Lower Austria and just one G in the plant from Italy. Downstream of this signal are 5 additional substitutions in the ETF of the two analyzed plants.

**Fig 5.**
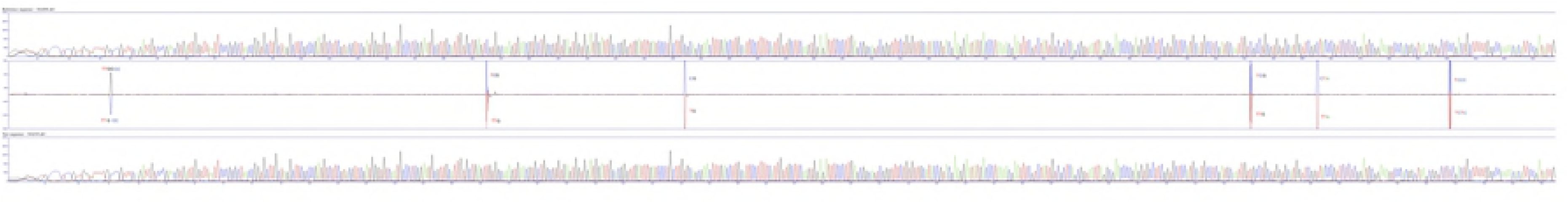
Alignment of two chromatograms. Alignment of nucleotide 100–500 of ETS sequence from a plant from Lower Austria (upper lane) and one plant from Cittiglio, Italy (lower lane). The lane in the middle is showing the difference with one additional G in the sequence of the Lower Austria region and five C/T substitution.

## Discussion

By sequencing the region of NTS/ETS/18S/ITS1/5.8S/ITS2 in the rDNA of *C. purpurascens* we could clearly distinguish the geographic region of the origin of the plant. So plants from Austria, Switzerland, Italy and France are having a unique signals in NTS/ETS, ITS1 and ITS2 (Fig 1, 3, 4, 5).

rDNA of plants from Austria get more diverse the farer we collect samples from Vienna. So plants from Styria and also from Pinzgau are having several substitutions in the analyzed region. Leaves collected in Lower and Upper Austria are very much conserved in the region of NTS/ETS/18S/ITS1/5.8S/ITS2. This conserved sequence could also be found in some plants from Salzburg, in a sample from Schaffhausen (Switzerland), Mühlheim an der Donau (Germany) and also from Orljavac (Croatia). Surprisingly it could not be found in the plants from Brigachtal (Germany) and Schleitheim (Schaffhausen, Switzerland).

The plants with the conserved sequence similar to the region of Vienna are all growing close to Benedictine gardens. So the leaf from Schaffhausen was collected in the garden of Allerheiligen, Mühlheim an der Donau is close to Beuron Archabbey and near Orljavac was a Benedictine abbey of Saint Michael.

The Benedictines had as early as in the 6th century medicinal herb gardens and distributed many plants across the Alps. Salzburg was and is the center of the Benedictines with the abbey of St. Peter. Alpine cyclamen were called “Wolfgang Erdäpferl” by the locals and was said to have a special effect on snakebite, migraine headaches and as a philter. The roots were sold to Pilgrims as salvation and fertility symbol [5]. So we suggest that at some point *C. purpurascens* had to be replanted in the area of Salzburg. We assume that for this reintroduction people used plants from Benedictine gardens. The plants from Monastery gardens seem to origin from the forests of Vienna.

In the forests of Schleitheim in the Canton of Schaffhausen is another occurrence of *C. purpurascens* which was described in 1989 [1]. The plants found in 1971 are very much related to plants in Switzerland. Unfortunately just beside this plants someone has planted some tubers from the Benedictine gardens. Although there are inflorescences it seems that no seeds germinated, not in Canton Schaffhausen and also not in Germany. That’s why there are just small patches of alpine cyclamen and they did not spread.

Interestingly plants in canton Schaffhausen are not very close related to the *C. purpurascens* growing in Brigachtal which is just 30 km across the border. The plants in Brigachtal seem to relate very much to plants from Austria. But it is possible that alpine cyclamen are naive to limestone in Southern Black Forest / Schaffhausen region. Another very interesting region would be the Tatra Mountains in the Czech Republic which should be analyzed in further studies. Also molecular cytogenetic mapping of 5S and 35S rDNA loci was not addressed in this study and should be considered in future studies.

## Conclusion

This is the first molecular analysis of the geographic origin of *C. purpurascens*.

An evolutionary and geographically interpretation in a phylogenetic context has been possible. rDNA sequences are variable within the species and some regions show a specific patten. With this information it is possible to determine the geographic origin of alpine cyclamen and to hypothesizing if it is native or was introduced at some point.

## Acknowledgements

The author would like to thank the members of the Flora Austria (Verein zur Erforschung der Flora Österreichs) particularly Hermann Falkner, Stefan Lefnaer, Georg Pflugbeil, Markus Sabor, Michael Strudl and Maria Zacherl for sending leaves of *C. purpurascens* from all over Austria. Without their precious help this work would not have been possible. Many thanks to Petra Bachmann and Peter Braig of the Canton Schaffhausen and Peter Enz, garden manager at the Zurich Botanical Garden for their help. Thanks to Duca Jaag for the plant from Orljavac. This research was supported by Thomas Kring.

The author received no specific funding for this work.

